# Egocentric belief attribution enables higher level Theory of Mind computation in group interactions

**DOI:** 10.64898/2026.03.27.714726

**Authors:** Shen Zhang, Huagen Wang, Raymundo Báez-Mendoza

## Abstract

Accurately attributing belief to others is a milestone of social cognition and serves as an indicator of intact cognitive functioning. However, tracking all others’ beliefs in complex social interactions is often inefficient or even impossible. We propose that in such cases, attributing one’s own belief to others (i.e., egocentric belief attribution) may serve as a functional approximation, enabling individuals to interact with others successfully. In this study, we employed computational modeling with behavioral measures to investigate this approach in social exchange in a novel group decision-making task across three experiments. We found that, beyond the well-documented reciprocity, participants exhibited consistent alternating behavior, characterized by the switching between potential recipients. This alternation was not driven by fairness concerns but reflected a strategic balance between maintaining stable partnerships and exploring alternatives. Crucially, a reinforcement learning model incorporating egocentric belief attribution (eToM) consistently outperformed all alternative models. These findings highlight that egocentric belief attribution is not a cognitive deficit, but a core mechanism supporting adaptive decision-making in complex social environments.

## Introduction

Sally was playing with her toy when her mother asked her to come over. She placed the toy into a basket and left. Later, her sister Anne saw the toy in the basket and assumed it had been forgotten, so she moved it to a drawer. When Sally returned, where would she look for her toy? Though the answer seems straightforward, children under the age of four often say that Sally would search the drawer. Correctly answering this question requires understanding that others can hold beliefs (here, a false belief) different from one’s own. Young children typically fail this task, attributing their own knowledge to others, a sign of egocentric thinking (Baron-Cohen et al., 1985; Scott & Baillargeon, 2017; Wimmer & Perner, 1983).

The ability to understand false belief by age four marks a key milestone in theory of mind (ToM) development, a foundational capacity for successful social interaction (Wellman et al., 2001). Children with autism spectrum disorder (ASD) consistently struggle with this task, failing to distinguish others’ beliefs from their own (Baron-Cohen et al., 1985; Senju, 2012). Similarly, egocentric belief attribution is observed in neurodegenerative conditions such as Alzheimer’s disease (AD, Lucena et al., 2020) and behavioral variant frontotemporal dementia (bvFTD, Gregory et al., 2002). These findings highlight that the capacity to attribute accurate mental states to others serves as a sensitive indicator of intact social cognition and broader cognitive health.

However, successful social interactions demand more than the basic false belief understanding assessed by the Sally-Anne task. One must track not only what A knows about B’s belief (i.e., a second-order belief), but also what anyone knows about any other’s belief. With more individuals involved in a social interaction, the number of possible belief states grows exponentially (Brandenburger & Dekel, 1993), making full tracking computationally inefficient. Indeed, even healthy adults often simplify processing by attributing their own beliefs to others in complex belief attribution task (Ereira et al., 2020), a phenomenon known as self-other mergence. More critically, in real-world social scenarios, actions are typically driven by multiple, intertwined beliefs that cannot be easily disentangled. For example, if both A and B invite C to dinner, it is often impossible to determine whether C accepted A’s invitation because A was perceived as kind or B as unkind from C’s experience. In such cases, attributing one’s own beliefs to others becomes not just a cognitive shortcut but a necessary strategy.

Here, we propose that, while often viewed as a sign of cognitive limitation, the egocentric belief attribution may instead serve as a functional approximation when tracking others’ beliefs becomes computationally infeasible. This perspective motivates a novel computational approach to modeling belief structures in repeated social interactions. Group interactions provide an ideal testbed, as individuals’ actions typically depend on their beliefs about multiple others—offering the rich social context necessary to probe such mechanisms. In the current study, we combine computational modeling with behavioral data from three experiments to investigate real-time group dynamics. We focus particularly on the role of egocentric belief attribution in shaping social decisions.

The Giving Game was used for this inquiry (Báez-Mendoza et al., 2021). Briefly, in this repeated game, on each trial, one of three players is randomly selected as the Actor. The actor is endowed with a token and must send it to one of the other two players. The reciprocity rule that people prefer to return other’s resource has been proposed as a key predictor of behavior in such social exchange settings. Consequently, in this game, an individual’s decision depends on their belief about how much each of the other two players has given them. This structure provides a natural context for testing the approach of egocentric belief attribution. The game was first implemented in Experiment 1. The results showed that participants tended to distribute their resources evenly and reciprocally among other players, establishing a behavioral baseline for modeling belief structures of Theory of Mind (ToM). After these results were established, we explored in Experiment 2 whether participants’ preference for even distribution stemmed from fairness concerns. Across both experiments, we compared simple reinforcement learning (RL) models with a ToM-based model incorporating egocentric belief attribution (eToM) and found that the e-ToM-based model provided a better fit to the data. This led to Experiment 3, in which we further examined in detail how the e-ToM model outperformed the other models, providing evidence for the involvement of e-ToM reasoning in such group-level social cognition.

## Results

### Experiment 1

In Experiment 1, 93 participants in 31 triads played a 90-trial repeated Giving Game. On each trial, one player was selected randomly as the Actor and sent one endowed token to one of the other two players, the Receivers (See Figure 1a and Methods). Following our previous study (Báez-Mendoza et al., 2021), we focused on how the decision maker’s last choice and the subsequent behavior of the other players influenced their current decision.

**Figure 1.**
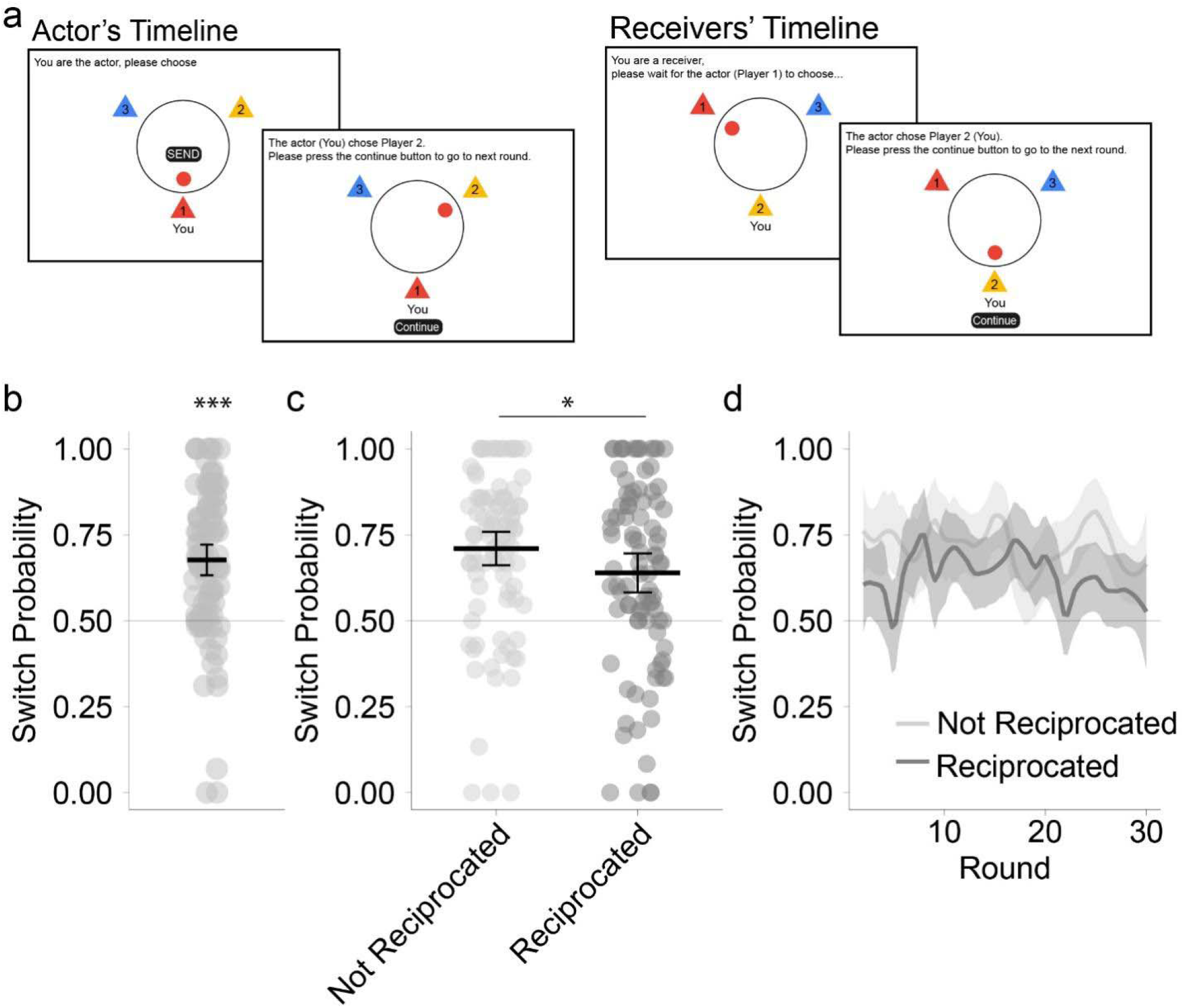
Experimental Design and model-free results of Experiment 1. **a)**Timeline of the Giving Game. On each trial, one player (the Actor) was pseudo-randomly assigned to send one endowed token to one of the other two players (the Receivers). The Actor role was assigned pseudo-randomly, ensuring that each player acted as Actor once per mini-block of three trials (one round) and never in two successive trials. **b)**Overall switch probability. Participants were more likely than chance to switch from one trial to the next which Receiver they gave a token to. **c)**Switch probability conditioned on whether the Actor was reciprocated. Switch probability decreased on trials when the Actor was reciprocated compared to trials when they were not. **d)**Switch probability did not change across rounds, and the linear trends in switch probability did not differ between trials when the Actors were reciprocated and trials when they were not. *indicates P < 0.05, **indicates P < 0.01, ***indicates P <0.001

Surprisingly, we found that participants were inclined to distribute their resources evenly across the other players. This pattern was reflected in a significantly higher than chance rate of alternating between recipients, in other words, the Actors switched who they offered a token from their previous decision to their current decision (Figure 1b, switch probability: mean = 67.64%, *β*(*SE*) = 1.95(0.27), *P <* 0.001; See method for analysis details). Moreover, as predicted, participants showed a tendency to reciprocate, that is, their switch probability decreased in trials where the rewarded player from their last choice returned at least one token back in response (the Actor was reciprocated in this case), compared to trials where no such reciprocation occurred (the Actor was not Reciprocated in this case; see Figure 1c, Not Reciprocated vs. Reciprocated: 70.96% vs. 63.87%, *β*(*SE*) = 0.40(0.15), *P =* 0.019).

After showing that participants favored both even distribution and reciprocal interactions, we further examined whether these tendencies evolved over time. However, switch probability did not change significantly across rounds (Figure 1d, *P =* 0.586), with no significant difference in the rate of change between trials when the Actor was reciprocated and whenthey were not (Figure 1d, *P =* 0.221).

Having established the participants’ behavioral patterns in the Giving Game, we sought to examine the computational mechanisms underlying these patterns. Specifically, we were to examine whether the eToM examine the data. We evaluated four candidate computational models (Figure 2b; See Methods for details); each capturing distinct aspects of the data. The first three models were motivated by the observed participants’ tendencies toward even distribution and reciprocity (See Methods for details). These three models assume that the players track all players’ sending behavior (including themselves) and make decisions based on their belief of their own sending behavior (The Alternating model), or the other two players’ sending behavior (The Reciprocity model), or both (The Alternating Reciprocity model). Specifically, the Alternating model assumes that the Actors send to the Receiver who received less from them previously. The Reciprocity model assumes the Actors send to the Receiver who gave them more in the past. The Alternating Reciprocity model integrates both mechanisms. In addition to these three heuristic models, we tested a fourth model incorporating an eToM component. This model extends the previous three by assuming the players not only track the others’ behavioral tendency, but also mentally represent their behavior as following the Alternating Reciprocity model. This mental representation enables the participants to adjust their estimation of others’ overall behavioral tendency. For example, after one player receives a token from an Actor, participants would infer that the tendency of this player to give to this Actor in the future would increase. Similarly, when one Actor gives a token to one Receiver, participants would infer that the Actor’s tendency to give to this Receiver in the future would decrease (given their preference for switching). Critically, as participants’ behavior are motivated both by their belief of their own actions and their belief of the others’ actions, these adaptation would not possible if no egocentric belief attribution is used (See methods). We refer to this model as the Mental Inference (Mental Infer) model reflecting its assumption that the players leverage a mental representation to infer others’ behavior. All models were fitted with a maximum likelihood estimation procedure (See Methods) and compared using the Akaike Information Criterion (AIC).

**Figure 2.**
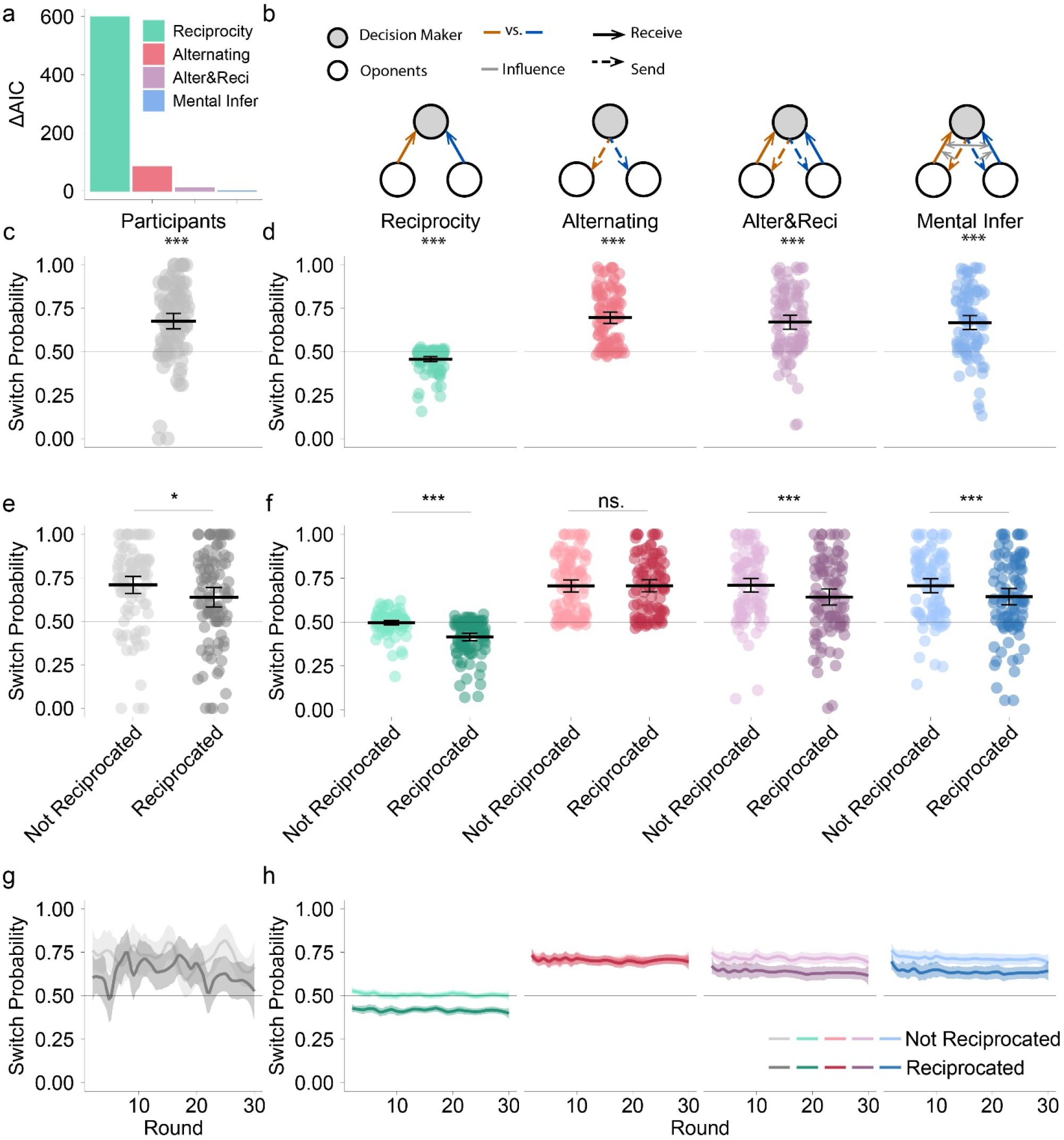
Model-based results of Experiment 1. **a)**Model fit comparison. The Mental Inference (Mental Infer) model achieved the lowest AIC, indicating the best fit to the data. **b)**Model assumptions. Four computational models were evaluated. The Reciprocity model assumes the Actors send to the Receiver who gave them more in previous rounds. The Alternating model assumes the Actors send to the Receiver who received less from them in previous rounds. And the Alternating Reciprocity model combines both mechanisms. The Mental Infer model further assumes players represent others’ behavior as following the Alternating Reciprocity model, enabling flexible adjustment of their estimation for others’ sending behavior. **c), d)** Switching behavior. c) Switch probability across trials copied from Figure 1b for comparison. d) All models except the Reciprocity model reproduced the above chance switch tendency. **e), f)** Reciprocity behavior. e) Switch probability decreased when the Actor was reciprocated, copied from Figure 1c for comparison. f) All models but the Alternating Model reproduced the reciprocity behavior of the participants. **g), h)** Temporal dynamics. g) No significant change in the switch probability was observed across rounds. And the linear trends of this switch probability change didn’t differ between trials when the actor was reciprocated and trials when the actor was not. However, h) While simulated data from all models showed no difference in the linear trends for the switch probability change between trials when the Actor was reciprocated and trials when the Actor was not, all models except the alternating model showed a slight but significant decrease in these switch probabilities. *indicates P < 0.05, **indicates P < 0.01, ***indicates P <0.001

Interestingly, the Mental Inference model outperformed all other models on the AIC metric (Figure 2a). To understand why, we conducted simulations with the best-fitted parameters from each model (See Methods) and computed the same measures as in the model-free analysis with the simulated datasets. This simulation analysis revealed that both the Alternating Reciprocity model and the Mental Inference model successfully captured the participants’ key behavioral patterns observed in the data. Specifically, **1) Switch probability:** All models but the Reciprocity model could reproduce the switching behavior (Figure 2d; switch probability vs. 50%; Reciprocity, M = 45.80%, *β*(*SE*) = −0.08(0.013), *P*<0.001; Alternating, M = 69.66%, *β*(*SE*) = 0.42(0.02), *P <* 0.001; Alternating Reciprocity, M = 67.03%, *β*(*SE*) = 0.35(0.03), *P <* 0.001; Mental Infer, M = 66.86%, *β*(*SE*) = 0.35(0.03), *P <* 0.001). **2) Reciprocity behavior:** All models but the Alternating model captured the reciprocity behavior shown by the relative decrease of switching probability on trials when the Actor was reciprocated (Figure 2f; Not Reciprocated vs. Reciprocated; Reciprocity, 49.58% vs. 41.44%, *β*(*SE*) = 0.09(0.01), *P <* 0.001; Alternating, 70.66% vs. 70.74%, *β*(*SE*) = 0.00(0.02), *P =* 0.973; Alternating Reciprocity, 71.03% vs. 64.33%,

β(*SE*) = 0.07(0.03), *P* =0.016; Mental Infer, 70.73% vs. 64.43%, *β*(*SE*) = 0.07(0.03), P =0.015). And **3) Temporal dynamics:** When examining the linear trends of the switch probability, no models showed a linear trend difference between trials when the Actor was reciprocated and trials when the Actor was not reciprocated (all *Ps* > 0.662). However, all models except the Alternating model exhibited a significant switch probability decrease over time (Reciprocity, *β*(*SE*) = −0.02(0.00), *P <* 0.001; Alternating, *β*(*SE*) = −0.00(0.00), *P =* 0.177; Alternating Reciprocity, 71.03% vs. 64.33%, *β*(*SE*) = −0.01(0.00), *P* <0.001; Mental Infer, *β*(*SE*) = −0.02(0.03), *P <* 0.001). Overall, Experiment 1 revealed that participants distributed their resources evenly and reciprocally across group members. These patterns were captured by both the Alternating Reciprocity and the Mental Infer model. However, no significant change in switch probability was observed across rounds, and all models except the Alternating model failed to capture this temporal stability. The overall above-chance switch probability is a finding that we didn’t expect. We were interested in whether such behavior was driven by fairness concerns. In the current design, however, players’ relative earnings were inherently tied to interaction dynamics, particularly reciprocity, making it difficult to disentangle fairness from strategic or adaptive behavior. To address this confound, we conducted Experiment 2 to isolate the role of fairness by manipulating the payoff structure and eliminating the correlation between earnings and reciprocity.

### Experiment 2

The above chance average switching rate observed in Experiment 1 was not predicted a priori, we therefore sought to replicate and examine this effect in Experiment 2. This pattern is consistent with well-documented fairness preferences in social decision-making. Thus, in addition to validating the robustness of the eToM mechanism in an independent cohort, a secondary aim of Experiment 2 was to test whether the switching behavior observed in Experiment 1 was driven by fairness concerns. To this end, we introduced a random deduction step to the Giving Game. Specifically, after the Actor sends the token, one token is deducted randomly from one of the Receivers’ accounts (Figure 3a). We refer to this modified game as the Giving Game with deduction. Because the deduction was assigned randomly, it was orthogonal to the players’ past actions and the game’s underlying dynamics. Thus, if the switching was driven by fairness concern, we expected a lower switching probability if the previously rewarded Receiver was deducted more tokens than the other player. In other words, we expected actors to favor the Recipient with greater recent losses. A new sample of 90 participants, organized into 30 triads, completed a 90-trial repeated Giving Game with Deduction.

**Figure 3.**
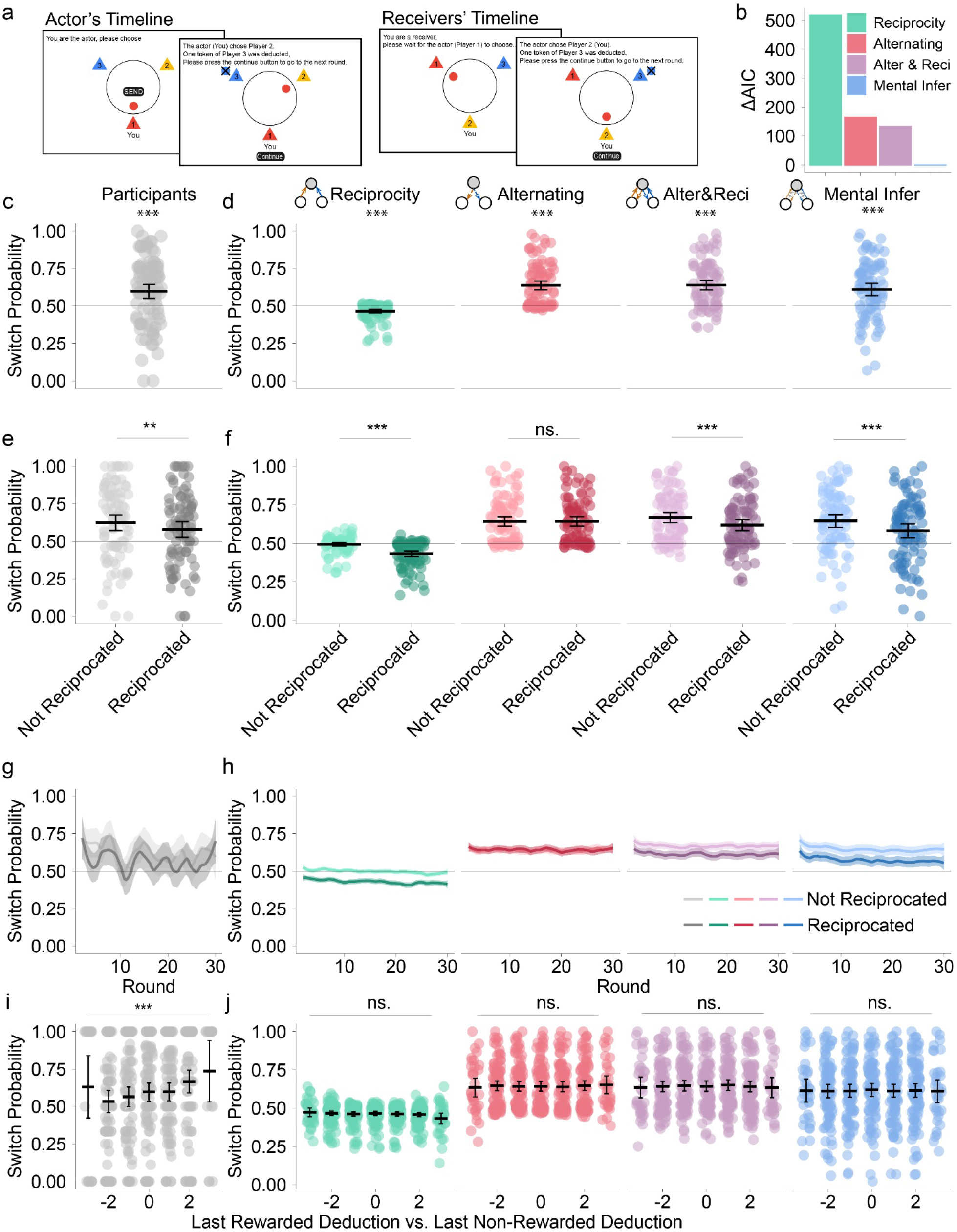
Experimental design and results of Experiment 2. **a)**Timeline of the Giving Game with deduction. On each trial, the Actor sends his/her token to one of the Receivers by rotating the table while the Receivers remain passive. After the Actor’s decision, one token is deducted from the account of one random Receiver. The results are then revealed to all players. **b)**Model fit comparison. The Mental Infer model achieved the lowest AIC, indicating the best fit of the data. **c) d)**Switching probability. c) The participants’ switch probability is significant above chance. d) All models but the Reciprocity model captured this switching tendency. **e), f)** Reciprocity behavior. c) Switch probability conditioned on whether the Actor was reciprocated. Switch probability decreased on trials when the Actor was reciprocated compared to trials when they was not. f) All models but the alternating model reproduced this pattern. **g), h)** Temporal dynamics. g) Switch probability decreased significantly across rounds, but this linear trend was not different between trials when the Actor was reciprocated and trials when the Actor was not. And, h) All but the Alternating model reproduced the decreasing trend in switch probability over time. **i), j)** Effect of deduction on switching. i) The greater the deduction from the previously rewarded Receiver relative to the other Receiver, the higher the probability of switching. j) None of the models captured this effect, as none explicitly incorporated the deduction mechanism. *indicates P < 0.05, **indicates P < 0.01, ***indicates P <0.001

We first conducted the same analyses as in Experiment 1 and found that the core behavioral patterns were largely replicated in this cohort. **1) Switch probability**. Again, participants distributed their resources evenly across the other players, as revealed by a significant above chance switching probability (Figure 3c; vs. 50%, M = 59.70%, *β*(*SE*) = 0.95(0.24), *P <* 0.001). All models except the Reciprocity model (Figure 3d; M = 46.46%, *β*(*SE*) = −0.07(0.01), *P <* 0.001) captured this pattern (Figure 3d; Alternating, M = 63.65%, *β*(*SE*) = 0.28(0.02), *P <* 0.001; Alternating Reciprocity, M = 63.82%, *β*(*SE*) = 0.28(0.02), *P <* 0.001; Mental Infer, M = 60.93%, *β*(*SE*) = 0.23(0.03), *P <* 0.001). **2) Reciprocity behavior**. Similarly, participants showed a tendency to reciprocate, indicated by a relatively lower switching probability when they were reciprocated (Figure 3e, Not Reciprocated vs. Reciprocated, 62.34% vs. 57.87%, *β*(*SE*) = 0.28(0.10), *P =* 0.009). This pattern was captured by all models except the Alternating model (Figure 3f; Reciprocity, 49.21% vs. 43.17%, *β*(*SE*) = 0.07(0.01), *P <* 0.001; Alternating, 64.19% vs. 64.31%, *β*(*SE*) = 0.00(0.02), *P =* 0.932; Alternating Reciprocity, 66.74% vs. 61.80%, *β*(*SE*) = 0.06(0.02), *P =* 0.023; Mental Infer, 64.47% vs. 58.19%, *β*(*SE*) = 0.07(0.03), *P =* 0.016). **3) Temporal dynamics**. Finally, we observed in this cohort that the switch probability decreased significantly across rounds, indicated by a negative linear trend of the switch probability with respect to the trial number (β(*SE*) = −0.29(0.13), *P =* 0.030). This trend still did not differ between trials when the Actor was reciprocated and the trials when he/she was not (β(*SE*) = −0.10(0.09), *P =* 0.287). All models but the Alternating model captured this temporal decline in switch probability (Reciprocity, *β*(*SE*) = −0.02(0.00), *P <* 0.001; Alternating, *β*(*SE*) = −0.00(0.00), *P =* 0.151; Alternating Reciprocity, *β*(*SE*) = −0.00(0.00), *P =* 0.004; Mental Infer, *β*(*SE*) = −0.03(0.00), *P <* 0.001).

After replicating the core behavioral patterns from Experiment 1, we examined the effect of the random deduction on participants’ decisions. We computed the difference in tokens deducted between the Receiver who received the token from the Actor’s last decision and the other Receiver and tested whether this difference predicted switch probability in a linear mixed model. Surprisingly, contrary to the fairness concern assumption, the more tokens deducted from the previously rewarded Receiver relative to the other Receiver, the higher the probability that the Actor switched their choice in the current trial. In other words, Actors were more likely to give the Receiver that had lost less tokens. This was revealed by a significant positive trend of the switch probability with respect to the deduction difference (Figure 3i, *β*(*SE*) = 0.15(0.04), *P <* 0.001). This effect was not captured by any of the computational models, as none explicitly accounted for such deduction (Figure 3j).

The Mental Infer model again outperformed all other models in terms of AIC (Figure 3b). However, the Alternating Reciprocity model also captured most of the key behavioral patterns observed in the data, matching the Mental Infer model in its ability to reproduce the switching tendency, Reciprocity behavior, and temporal dynamics. This raises a critical question: do participants really have a mental model? This question is central because only if individuals represent others’ mental states can belief attribution, such as egocentric reasoning be considered a genuine cognitive process. To address this, we designed Experiment 3 to directly test whether participants maintain a mental model of others in social exchange.

### Experiment 3

The aim of Experiment 3 was to further examine whether the participants actually maintain a mental representation of others. To probe this, we adapted the game structure from Experiment 1: during the Actor’s decision, the Receivers are asked to guess whom the Actor will choose (Figure 4a; see Methods for details). Critically, these guesses have no impact on all players’ earnings. This design ensured that the payoff structure in Experiment 3 matched that of Experiment 1, minimizing any influence of the guessing task on participants’ behavior. We refer to this modified game as the Giving Game with Guesses.

**Figure 4.**
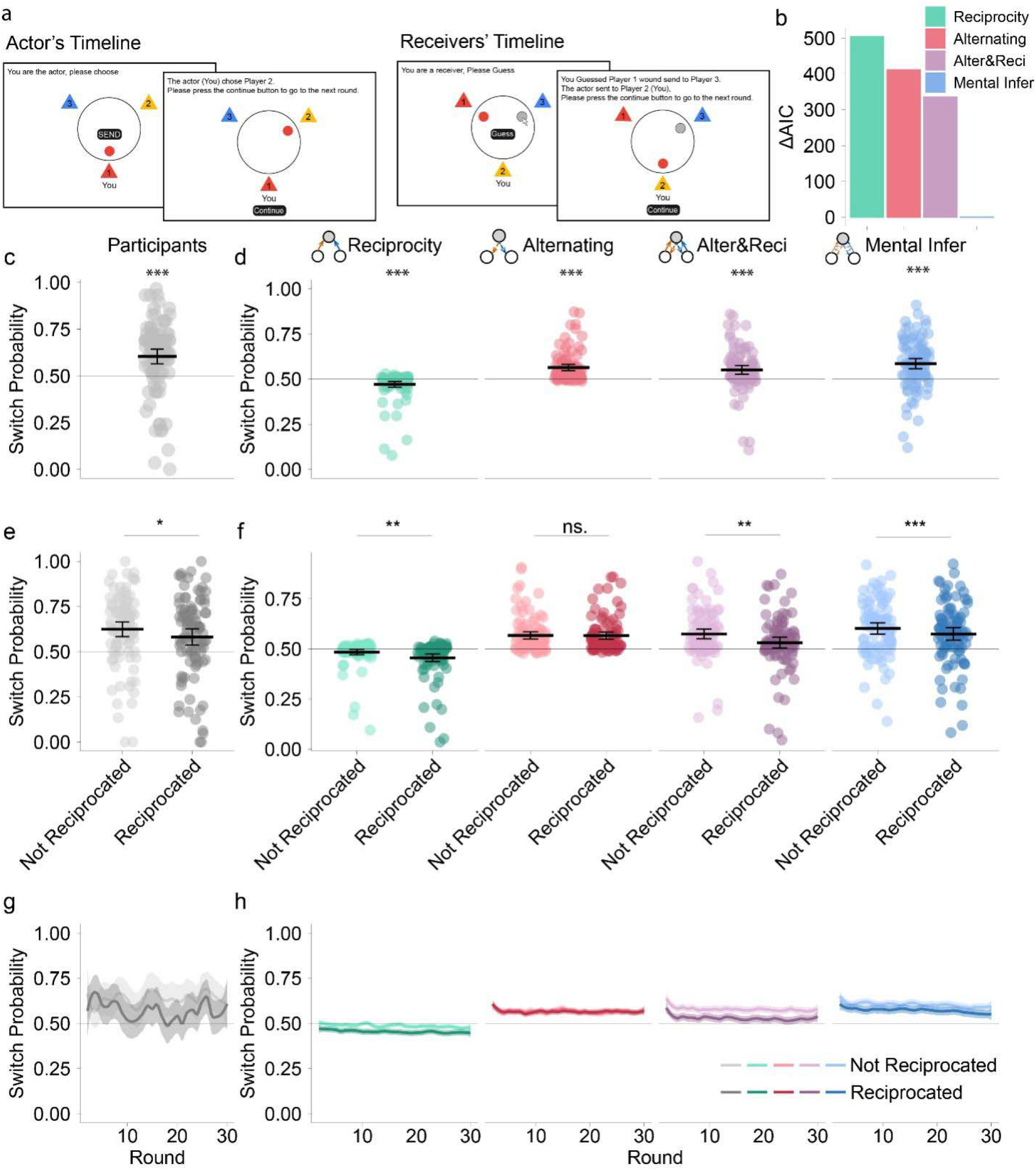
Experimental design and results of sending behavior in Experiment 3. **a)**Timeline of the Giving Game with guesses. On each trial, one Actor is randomly selected to send one token to one of the other players (the Receivers). Simultaneously, the Receivers guess whom the Actor will choose. After all decision are made, only the Actor’s decision is revealed to all players. **b)**Model fit comparison. The Mental Inference (Mental Infer) model achieved the lowest AIC, indicating the best fit to the data. **c), d)** Switch probability. c) The participants’ switch probability was significantly above chance. d) All models except the Reciprocity model reproduced this switching tendency. **e), f)** Reciprocity behavior. e) The switch probability decreased when the Actor was reciprocated. f) All models except the Alternating model reproduced this behavior pattern was captured. **g), h)** Temporal dynamics. g) The switch probability did not change significantly across rounds, and the linear trend did not differ between trials when the actor was reciprocated and trials wen the actor was not. h) All models reproduced a small but significant decrease in switch probability over time, and showed no difference in this linear trends between trials when the Actor was reciprocated and trials when the Actor was not, *indicates P < 0.05, **indicates P < 0.01, ***indicates P <0.001

We tested a 90-trial Giving Game with Guesses in a cohort of 96 participants in 32 triads. First, we performed the same model-free analyses as in Experiment 1. Importantly, when performing the model-based analyses, apart from the sending choices, the participants’ guessing choices were also fitted by all models. Particularly, for the three naïve models, the Receivers’ guesses of the Actors’ choice were only based on the estimation of the Actors’ sending tendency, as no mental representation was assumed. Adopting these models makes the participants’ participants’ probability of guessing other’s switch below chance. In contrast, the Mental Infer model allowed the participants to use a mental representation to guide their guesses, by simulating others’ choices according to the Alternative Reciprocity model (See Methods for details). This mental simulation would make the participants’ probability of guessing other’s switch significantly above chance.

Similar behavioral patterns were observed in Experiment 3, and the Mental Infer model again achieved the best fit, as indicated by the lowest AIC (Figure 4b). Particularly, **1) Switch probability**. Participants distributed their own resources evenly across other players, evidenced by an significant above-chance switching probability (Figure 4c; vs. 50%, M = 60.34%, *β*(*SE*) = 0.93(0.19), *P <* 0.001). All models except the Reciprocity model captured this tendency (Figure 4d; Reciprocity, M = 47.10%, *β*(*SE*) = −0.06(0.01), *P <* 0.001; Alternating, M = 56.41%, *β*(*SE*) = 0.13(0.01), *P <* 0.001; Alternating Reciprocity, M = 55.13%, *β*(*SE*) = 0.11(0.01), *P <* 0.001; Mental Infer, M = 58.51%, *β*(*SE*) = 0.17(0.03), *P <* 0.001). **2) Reciprocity behavior**. Participants also showed a tendency to reciprocate, indicated by a relatively lower switching probability when they were reciprocated (Figure 4e, Not Reciprocated vs. Reciprocated; 62.37% vs. 58.13%, *β*(*SE*) = 0.21(0.10), *P =* 0.038). This pattern was captured by all models except the Alternating model (Figure 4f; Reciprocity, 48.38% vs. 45.53%, *β*(*SE*) = 0.03(0.01), *P =* 0.005; Alternating, 56.74% vs. 56.75%, *β*(*SE*) = 0.00(0.01), *P =* 0.909; Alternating Reciprocity, 57.45% vs. 53.08%, *β*(*SE*) = 0.05(0.02), *P =* 0.007; Mental Infer, 60.15% vs.57.39%, *β*(*SE*) = 0.03(0.01), *P <* 0.001). 3) Temporal dynamics. Finally, switch probability did not change significantly across rounds, indicated by a non-significant linear trend of the switch probability with respect to the round number (β(*SE*) = 0.00(0.11), *P =* 1.00). And this trend also didn’t differ between trials when the Actor was reciprocated and those when he/shewas not (β(*SE*) = 0.10(0.09), *P =* 0.265). However, all models produced a negative linear trend in switch probability changes (Reciprocity, *β*(*SE*) = −0.01(0.00), *P <* 0.001; Alternating, *β*(*SE*) = −0.00(0.00), *P =* 0.007; Alternating Reciprocity, *β*(*SE*) = −0.01(0.00), *P <* 0.001; Mental Infer, *β*(*SE*) = −0.02(0.00), *P <* 0.001). These findings replicated the core patterns observed in Experiment 1, confirming the robustness of the behavioral dynamics across conditions.

After showing a similar pattern in participants’ sending behavior, we then examined whether they could correctly guess others’ sending behavior. Overall, the participants were able to predict the Actors’ choices above chance level, as evidenced by a guess accuracy significantly above 50% (Figure 5a, vs. 50%, M = 54.76%, *β*(*SE*) = 0.41(0.09), *P <* 0.001). This pattern was only captured by the Mental Infer model (Figure 5b; M = 53.97%, *β*(*SE*) = 0.08(0.01), *P <* 0.001) and by the Reciprocity Model (M = 51.73%, *β*(*SE*) = 0.04(0.01), *P <* 0.001) but not by the other two models (Alternating, M = 49.30%, *β*(*SE*) = −0.02(0.00), *P <* 0.001; Alternating Reciprocity, M = 50.53%, *β*(*SE*) = 0.01(0.01), *P =* 0.20). The guess accuracy did not differ between trials when the Actor was reciprocated and trials when the Actor was not (Figure 5c, Not Reciprocated vs. Reciprocated; 53.33% vs. 57.36%, *β*(*SE*) = −0.12(0.07), *P =* 0.103), showing similar performance in these two kinds of trials. All models reproduced this pattern (Figure 5d; Reciprocity, 51.52% vs. 52.01%, *β*(*SE*) = −0.00(0.01), *P =* 0.533; Alternating, 49.08% vs. 49.20%, *β*(*SE*) = −0.00(0.00), *P =* 0.554; Alternating Reciprocity, 50.22% vs. 50.75%, *β*(*SE*) = −0.01(0.01), P =0.489; Mental Infer, 54.11% vs. 54.08%, *β*(*SE*) = 0.00(0.01), *P =* 0.973). More interestingly, the participants improved their performance over time, indicated by a small but significant positive linear trend in the guess accuracy (Figure 5e; *β*(*SE*) = 0.15(0.07), *P =* 0.024).

**Figure 5.**
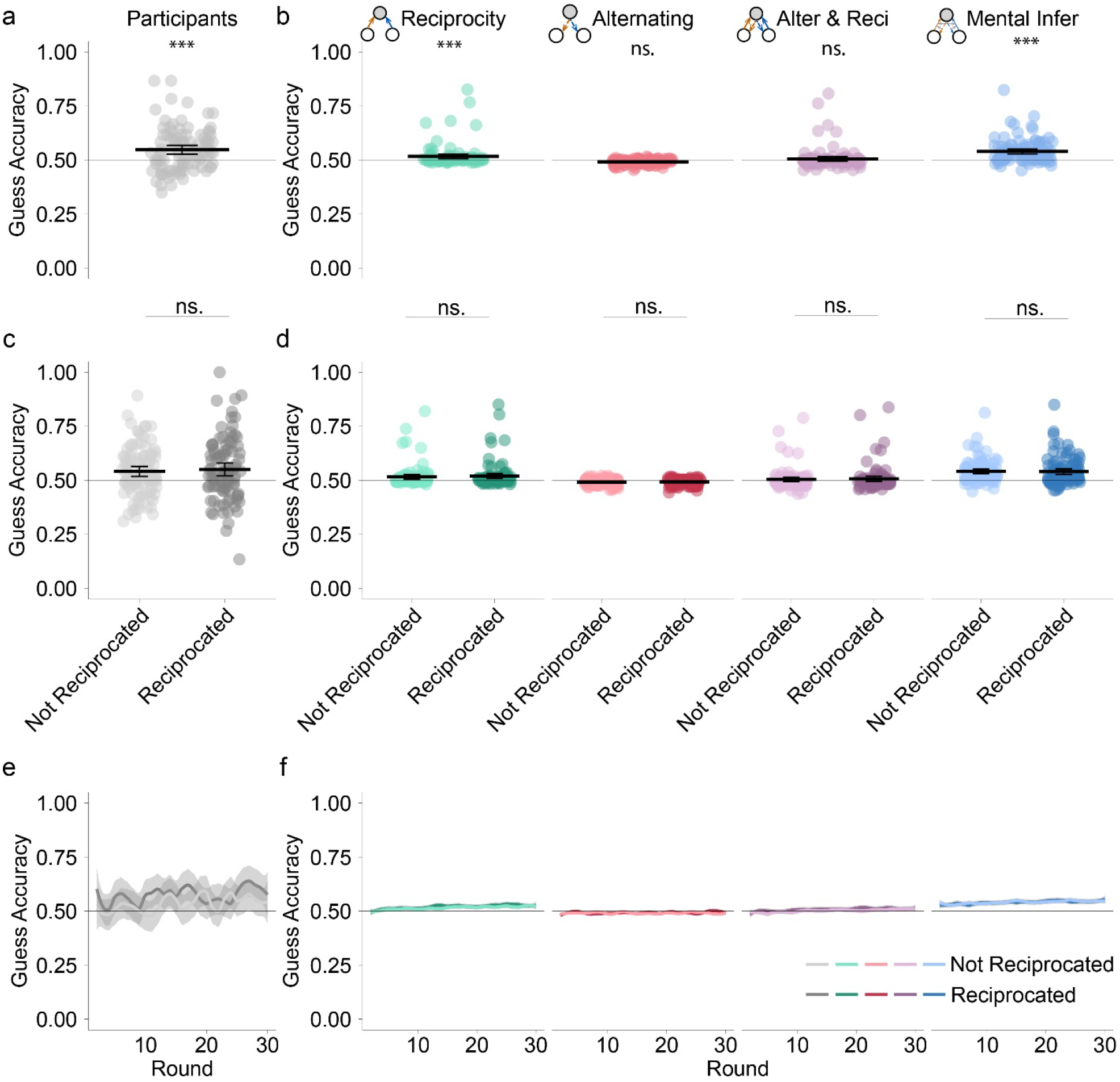
Guessing accuracy in Experiment 3. **a), b)** Guessing accuracy. a) The participants’ guessing accuracy was above chance. b) Only the Mental Infer model and the Reciprocity model captured this above-chance accuracy. **c), d)** Guessing accuracy conditioned on whether the Actor was reciprocated. c) The guessing accuracy did not differ between trials when the Actor was reciprocated and the trials when the Actor was not, and d)All models reproduced the effect in c). **e), f)** Improvement in guessing accuracy. e) Participants’ guessing accuracy increased across rounds. f) All models except the Alternating model reproduced the temporal improvement in guessing accuracy. *indicates P < 0.05, **indicates P < 0.01, ***indicates P <0.001

**Figure 6.**
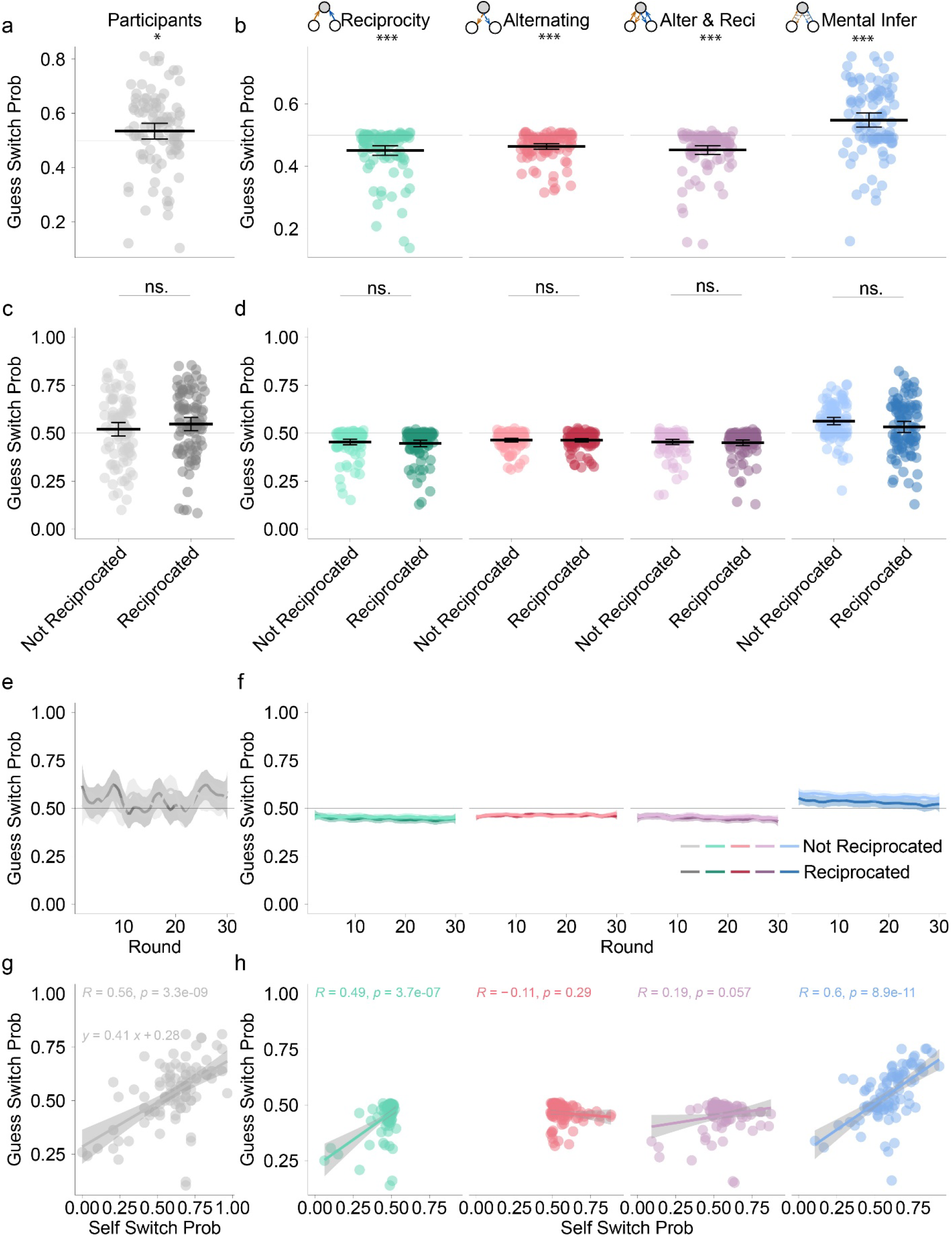
Behavior of guessing a switch in Experiment 3. **a), b)** Probability of guessing a switch. a) Participants are more likely than chance to guess the Actor would switch choices. **b)** Only the Mental Infer model captured the above-chance probability of guessing a switch. **c), d)** Probability of guessing a switch conditioned on whether the Actor was reciprocated. c) The probability of guessing another switch didn’t differ between trials when the Actor was reciprocated and the trials when the Actor was not. d) All models reproduced the effect. **e), f)** Changes of probability of guessing a switch over time. e) Participants’ probability of guessing a switch did not change across rounds. f) The Reciprocity and the Alternating model captured this effect. **g), h)** Alignment of self and guess switching behavior. g) The participants’ guessing probability of whether the Actor would switch was correlated with their own switch probability. h) This behavioral alignment was only reproduced by the Mental Infer model and the Reciprocity model. *indicates P < 0.05, **indicates P < 0.01, ***indicates P <0.001

All models except the Alternating model captured this liner trend (Figure 5f; Reciprocity, *β*(*SE*) = 0.01(0.00), *P* < 0.001; Alternating, *β*(*SE*) = −0.00(0.00), *P =* 0.139; Alternating Reciprocity, *β*(*SE*) = 0.01(0.00), *P <* 0.001; Mental Infer, *β*(*SE*) = 0.01(0.00), *P =* 0.002). This linear trend didn’t differ between the trials when the Actor was reciprocated and the trials when the Actor was not (Figure 5e; Not Reciprocated vs. Reciprocated; *β*(*SE*) = 0.00(0.06), *P =* 0.935). All models reproduced this effect (Figure 5f; all Ps > 0.156).

Having established that the participants’ guessing accuracy was significantly above chance, a pattern which can be only captured by the Mental Infer model, we finally examined what guess patterns may lead to such above chance accuracy. Specifically, we examined whether the participants anticipated the Actor would switch their choices, and whether this expectation was adapted through the interaction process. We found that participants believed that others would distribute their resources evenly, i.e., they guessed that the Actors would switch their choices. This was demonstrated by participants’ above-chance performance in guessing an option different from the Actor’s last choice (Figure 5g; vs. 50%, M = 53.47%, *β*(*SE*) = 0.29(0.13), *P =* 0.027). This pattern was produced only by the Mental Infer model (Reciprocity, M = 45.15%, *β*(*SE*) = −0.10(0.02), *P <* 0.001; Alternating, M = 47.48%, *β*(*SE*) = −0.07(0.01), *P <* 0.001; Alternating Reciprocity, M = 45.23%, *β*(*SE*) = −0.10(0.02), *P <* 0.001; Mental Infer, M = 54.80%, *β*(*SE*) = 0.10(0.02), *P <* 0.001). We next tested whether the participants expected the Actors to reciprocate with others. We found no evidence for this: the probability of guessing a switch did not differ between trials when the Actor was reciprocated and trials when s/he was not (Figure 5i; Not Reciprocated vs. Reciprocated, 52.11% vs. 54.71%, *β*(*SE*) = 0.06(0.09), *P =* 0.486). All models reproduced this pattern (Figure 5j; Reciprocity, 46.41% vs. 44.96%, *β*(*SE*) = 0.01(0.02), *P =* 0.583; Alternating, 49.85% vs. 49.74%, *β*(*SE*) = 0.00(0.01), *P =* 0.912; Alternating Reciprocity, 49.46% vs. 47.70%, *β*(*SE*) = 0.01(0.02), *P =* 0.912; Mental Infer, 57.60% vs. 54.65%, *β*(*SE*) = 0.03(0.02), *P =* 0.189). We then examined whether the participants’ probability of guessing a switch changed across rounds. No significant linear trends in probability of guessing a switch were observed (Figure 5k; *β*(*SE*) = 0.06(0.07), *P =* 0.399). However, the Alternating Reciprocity model and the Mental Infer model reproduced a significant negative linear trend, while the other models didn’t (Figure 5l; Reciprocity, *β*(*SE*) = −0.01(0.00), *P =* 0.703; Alternating, *β*(*SE*) = 0.00(0.00), *P =* 0.916; Alternating Reciprocity, *β*(*SE*) = −0.01(0.00), *P =* 0.042; Mental Infer, *β*(*SE*) = −0.02(0.00), *P =* 0.002). This linear trend didn’t differ between the trials when the Actor was reciprocated and the trials when the Actor was not (Figure 5k; Not Reciprocated vs. Reciprocated; *β*(*SE*) = 0.04(0.06), *P =* 0.496). All models reproduced this effect (Figure 5l; all Ps > 0.506). Finally, we examined whether the participants’ mental representations were aligned with their own behavior. Interestingly, we found that the participants who distributed their resources more evenly would also think others would do the same. That is, participants own switching probability was correlated with their guesses of others’ switching probability (Figure 5m, R=0.56, P<0.001). This alignment was captured by the Mental Infer model (Figure 5n; R=0.60, P<0.001) and the Reciprocity model (R=0.49, P< 0.001), but not the other two models (Alternating, R = −0.11, *P =* 0.290; Alternating Reciprocity, R = 0.15, *P =* 0.057).

## Discussion

Attributing accurate mental states to others has long been regarded as a hallmark of intact cognitive functioning. In the current study, we proposed that in complex social interactions, attributing one’s own beliefs to others (i.e., the egocentric belief attribution) may serve as a necessary approximation when tracking all others’ beliefs becomes computationally infeasible. This hypothesis was tested using a novel social exchange game, the Giving Game, combined with computational models. We showed that participants distributed their resources evenly and reciprocally among other players in repeated giving games. This tendency toward equal distribution was not primarily driven by a concern for fairness as in accrued rewards, but rather by a preference for allocating resources equitatively. Furthermore, the computational model that incorporated the egocentric belief attribution (i.e., the Mental Inference model) consistently outperformed all other models under consideration. This suggests that ToM plays a critical role in guiding resource-sharing decisions, enabling individuals to anticipate others’ behavior and adapt their own choices accordingly.

It is well documented that when learning about others’ preferences, individuals initially use their own preferences as a reference point (Zhang et al., 2025). Our computational approach aligns with this phenomenon and extends it to the domain of high-level belief attribution in complex social interactions. Together with this, our modeling results suggest that egocentric belief attribution may be a general approach the cognitive system adopted when information or cognitive resource is limited to support tracking of all possible mental states. This idea is further supported by evidence showing that when individuals attempt to remember information about both themselves and others, the two types of memory become intertwined, a phenomenon known as self-other mergency (Wittmann et al., 2016) a phenomenon known as serf-other mergence. Our results indicate that under conditions when accurate attribution is not possible, egocentric beliefs may dominate or override other, more accurate representations of others’ mental states.

The average switching probability was significantly above chance across all experiments. Given that fairness preference has long been documented to drive distributional behavior (Fehr & Schmidt, 1999; Sanfey et al., 2003), one might interpret the observed alternating behavior as a manifestation of fairness concerns. However, we manipulated the losses of the Receivers in Experiment 2 and found that, contrary to the prediction of fairness concern, the Actor’s switch behavior was positively associated with the relative deduction of the Receiver who received the token from the Actor’s last choice. This result suggests that the Actor’s alternating behavior was not due to the participants’ fairness preference per se.

We also found that when an Actor was reciprocated by the rewarded Receiver, their switching probability on the next choice would decrease. This pattern is in line with the prediction of the reciprocity rule proposed by social exchange theory (Cropanzano & Mitchell, 2005). Reciprocity has long been regarded as critical for cooperation (Nowak, 2006), enhancing group welfare (Fehr & Schmidt, 1999; Rand et al., 2009), and thus is conserved through evolution. Indeed, it has been proposed that people innately prefer reciprocity (Fehr et al., 2002). However, the reciprocity observed in our experiments cannot be fully attributed to the participants’ preference for reciprocity. As for most of the repeated games, following the reciprocity rule may help build a good reputation (Milinski et al., 2002), leading to higher long-term payoffs. Thus, participants’ reciprocating with others may also arise from a concern for reputation, rather than an intrinsic preference.

A decrease in switching probability over time was observed in Experiment 2, but not in Experiments 1 or 3. This temporal trend may reflect the formation of stable dyadic interactions in Experiment 2, which was absent in the other conditions. According to social exchange theory, the risk associated with resource sharing is positively correlated with the strength of the resulting social bond (Kollock, 1994; Molm et al., 2000; Savage & Whitham, 2025). In Experiment 2, we introduced a random deduction mechanism, creating a riskier environment than in the other experiments. This increased environmental risk may explain why a declining switching rate, indicative of relationship stabilization, was only observed in Experiment 2.

RL models have been successfully used to capture the dynamics in learning others’ behavioral tendencies, such as generosity (Hackel et al., 2015), willingness to cooperate (Jin et al., 2023), and fairness preference (Zhang et al., 2025). This enables us to model the first-level beliefs of the participants. Social exchange in groups requires tracking the behavioral histories of all group members, for example, who has given more in the past, suggesting that simple RL models can efficiently capture the core dynamics of group-level resource allocation. The computational models we evaluated captured the participants’ dual tendencies to reciprocate and to alternate. That is, individuals appeared to balance the trade-off between maintaining a stable partner and exploring alternative ones. Most prior studies have focused on how the reciprocity rule shapes the interaction outcomes, often overlooking the exploratory phase at the beginning of the social exchange trajectory (e.g., Karpiński et al., 2023). Our findings reveal that alternating behavior is a prominent feature of repeated interactions within groups, suggesting that decisions are shaped not only by reciprocity but also by an interplay with alternation. These results highlight the importance of examining the entire interaction trajectory of resource-sharing behavior in groups and the potential influence of multiple, often-overlooked factors that extend beyond the scope of traditional social exchange theories.

Successful social interactions require not only inferring others’ behavioral tendencies but also anticipating the impact of their own behavior on others. Modeling such hierarchy of beliefs has been achieved through interactive Partially Observable Markov Decision Processes (iPOMDP; Gmytrasiewicz & Doshi, 2005), game theory of mind (Yoshida et al., 2008), and meta-Bayesian models (Devaine et al., 2014). While successful in capturing individuals’ behavior in social interactions, all these models are computational demanding. A seminal study introduced a simplified h-ToM framework with a two level belief hierarchy and a delta rule based learning (Hampton et al., 2008). The Mental Infer model in the current paper extends this approach to the group settings where individuals’ decisions are driven by more than one belief. This approach incorporates a more complex mental model of others’ preferences while maintaining computational tractability. It’s critical that without the egocentric belief attribution, this extension would not be possible.

In Experiments 1 and 2, the Mental Infer model outperformed all the other models, yet the Alternating Reciprocity model still captured most of the participants’ behavior. However, in Experiment 3, all the basic reinforcement learning models failed to predict the participants’ guess behavior, highlighting the limitation of such models that lack a mental representation. This divergence further supports the validity of the Mental Infer model and suggests that individuals leverage their mental representation when allocating resources. To investigate whether this mental representation is learned over time or present prior to, we examined the guess behavior dynamics across all trials. Interestingly, even though the guessing accuracy increased over time, no significant change in the probability of guessing if others would switch was observed from early to late stages of the task. Together with the result that the switching probability of one’s own is positively correlated with their guess of others’, this also suggests that participants projected their own preference onto others as a reference in social interactions.

In summary, we tested a computational model of hierarchical belief representation in group interactions, grounded in egocentric belief attribution. We found that the reciprocity rule plays a key role in shaping resource-sharing behavior within groups. Moreover, participants frequently switched between potential recipients, highlighting the adaptive value of exploring alternative partners during social interactions. Crucially, the model incorporating egocentric belief attribution (the Mental Inference model) provided the best fit to behavioral data across all experiments. This suggests that egocentric reasoning is not a cognitive limitation, but a functional mechanism that supports hierarchical belief representation under conditions of cognitive constraint.

## Methods

### Participants

For Experiment 1 and Experiment 2, we recruited Europe-based participants from the online recruiting platform Prolific. We applied the IP address filter provided by Prolific to include participants from all European Union countries. However, almost all participants were from the UK. Groups that accomplished fewer than 20 trials were removed from the analyses and the data collection were stopped when the sample size of 30 groups was reached (Experiment 1: n = 93, Age (SD) = 36.62 (11.55), 58 females; Experiment 2: n = 90, Age (Sd) = 37.72 (11.30), 52 females) in both experiments. The experiments were approved by the Ethics Committee of the Georg-Elias-Müller Institute of Psychology of the University of Göttingen.

For Experiment 3, we recruited participants (n = 96, Age (SD) = 19.98 (1.86), 72 females) from the Beijing Forestry University, as no groups dropped out during the procedure, we did not remove any groups. Participants provided informed consent in accordance with the Beijing Forestry University, Research Ethics Board.

### Procedure

Experiments 1 and 2 started with a consent form explaining the study’s information (See A1 for the consent form). After reading the consent form, the participants needed to decide whether they want to participate in the study. The experiments only proceeded if they chose to take part in the study.

On the first page, the participants filled out two demographic questions (Age and Gender, see A1 for details). Then they were presented with the task and game interface instructions followed by some practice to get familiar with the interface (See A1 for the instructions). The participants could practice as many trials as they wanted and press a button to enter the waiting room in which they would be grouped with others. As soon as enough participants joined the waiting room, the program would arrange the participants into groups of three players and the main task would start. The participants will wait for at most 10 minutes in the waiting room and will be compensated with £1.50 if not enough participants enter the room during this period. As long as the task started, the participants would be compensated with £2 plus a bonus (in the Giving Game).

The same procedure was applied for experiment 3, except that participants came to the testing room to complete the task, and provided a paper form consent form. Experiment 3 took place in a testing room with several PCs. Participants in each group came to the testing room at the same time. They were asked not to talk to each other during the whole experiment. However, as they could see each other, we only recruited participants of the same gender for each group to control for potential gender influence on reciprocity behavior. The payment for the participants from experiment 3 is described below.

### The Giving Games

We adopted the Giving Game (Báez-Mendoza et al., 2021) to explore the reciprocity behavior in three experiments (see Figure 1). In all experiments, participants played the game in a group of three players. On any given trial, one player was pseudo-randomly selected as the Actor, and was endowed with a token. The Actor had to give this token to one of the other two players (the Receivers). After showing the Actor’s decision to all players, the trial ended, and all players pressed a Continue button to enter the next trial. We pseudo-randomized who was the Actor following these two rules: A player could not be the Actor two rounds in a row, and all players had to be in the role of Actor at least once every five trials, resulting in mini-blocks of three trials in which all participants were in the Actor role. The participants performed a maximum of 90 trials using the interface described in the Results section. Furthermore, if any of the participants failed to respond within 5 minutes on any trial, the task ended automatically, and the participants advanced to the debriefing page.

All participants who started the task were compensated with a base payment of £2 plus a bonus determined by the tokens they earned in the task, with a conversion ratio of 1:0.03 (tokens to £).

For Experiment 2, the game was the same as in Experiment 1, except that on each trial, a token was deducted from a randomly chosen Receiver. This deduction was revealed to all players together with the Actor’s decision. This setting allows us to test whether the alternating behavior we found in Experiment 1 was due to pure fairness or strategic concerns. We also gave an initial bonus of £0.30 to compensate for the potential losses during the game because of this random deduction procedure, and the conversion rate of the token to real money is still 1:0.03 (tokens to

£).

In Experiment 3, we wanted to test whether the belief about other players influences one’s own behavior. To do so, we had the Receivers predict whom the Actor would send to while the Actor was deciding. After all player made their decisions, the Actor’s decision was shown to all players, while the guesses of the Receivers were not. This setting allows the participants to update their guess according to the Actor’s behavior while removing the possibility of the guess behavior influencing others’ behavior. All players were compensated with a show-up fee of 15 CNY, and each token earned in the game was converted to 0.5 CNY.

### Model free analyses: the linear mixed models for behavior

We performed linear mixed model analyses for all dependent variables. In particular, the switch choices were modeled by fixed effects of whether the Actor was reciprocated, its interaction with trial number and random effects with the same terms of all fixed effects. The “whether reciprocated” factor was coded in two columns thus suppressing the intercept term. Its interaction with trial number was coded similarly. For the random-effects structure, we retained the maximal set of terms that allowed model convergence. The switch choices were coded as 1 if the Actor’s choice was different from their previous choice, 0 otherwise. The full specification is:

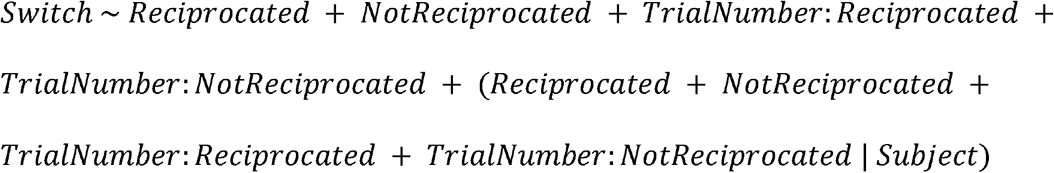

When considering other variables, the formulas are the same except the dependent variables. For guess accuracy, the choices were coded as 1 if the guess is correct, 0 otherwise. The guess switch choices were coded as 1 if the Receiver’s guess is different from the Actor’s previous choice, 0 otherwise. Finally in Experiment 2, the relative number of deduction are added both as a fixed and a random effect of the model. All liner mixed models are conducted using the lme4 package (Bates et al., 2015) under the R environment.

### The naive models: considering alternating and reciprocity

#### Notations

We denoted the three players as player *i* (*i* ∈ *N* = {1,2,3}). The player sitting at the counter-clockwise (ccw for short) side of player *i* is player *k*(*k* = *i* + 1, if *i* = 3, *k* = 1). This defines a *ccw* function mapping player *i* to its counter-clockwise side player. The function *cw* for *i*’s clockwise side player is defined similarly.

In this task, the primary goal of the players is to estimate the tendency (i.e., the probability) of player *i*’s giving to player *j*, denoted as *P*_*ij*_. Thus, *P*_12_ = 1 − *P*_13_ which motivated us to define *p*_*i*_ = *P*_*i*[*ccw*(*i*)]_ (i.e., *p*_1_ = *p*_12_). This leads to *P*_*i*[*cw*(*i*)]_ = 1 − *p*_*i*_. The estimate of player 1 on *p*_*i*_(*i,j* ∈ *N*) is denoted as 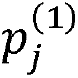. All models assume the players make decisions based on these estimates.

#### Model assumptions

For simplicity, we shall describe all computational models from the perspective of player 1. And with the help of our *ccw* and *cw* functions, all the models can be easily deduced from the perspectives of other players.

In the naive models, we assume the players have two main concerns.

#### Alternating

They distributed their resources evenly, i.e., the Actors tend to send their tokens to the Receiver who receive less tokens from their previous decisions. Put formally, player 1 tends to give to player 2 if 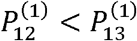, that is, 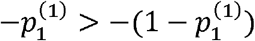 This term is to capture the switching behavior.

#### Reciprocity

The players are also motivated by the exchange history between them and other players. Specifcally, they tend to send to Receiver who gave them more in the previous trials. That is, player 1 gives to player 2 not player 3 if 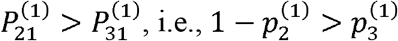

#### Updating estimate

All players update their estimates in a trial-by-trial manner. We have the following definitions:

Denote the actor of trial *t* as *a*(*t*), a function mapping integer number to *N*, and similarly, denote *r*(*t*) as the receiver who received the token on trial t.

Denote *R*_*i*_(*t*) as the reward *i* gave to *ccw(i)*, so we have:

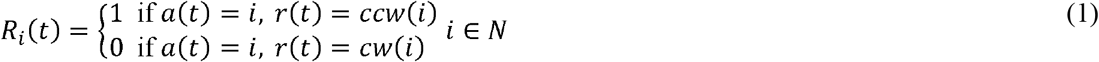

We then have the following update rules for *p*_*i*_ (player 1’s estimate):

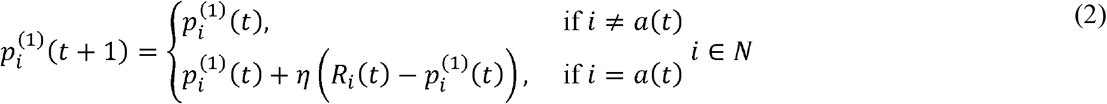

in which, *η* is the learning rate for this update.

#### Making decisions

All naive models differ only in the decision-making stage. The Alternating Reciprocity Model assumes the player makes decisions according to the other two players’ value to them. Formally, we denoted the value of player *i* to player *j(i ≠ j)* as 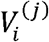, and it has two components:

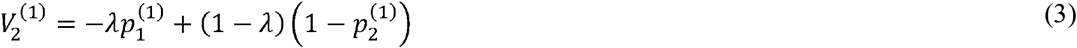

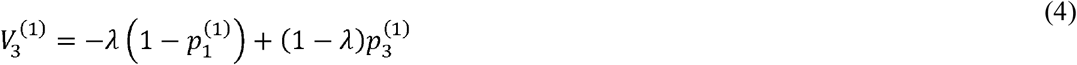

We omit the index of the trial number in the equation to keep it simple, this value is updated trial by trial. *λ* is the weight of each component for player 1. The probability of sending to player 2 when player 1 is the actor is given by the Luce rule:

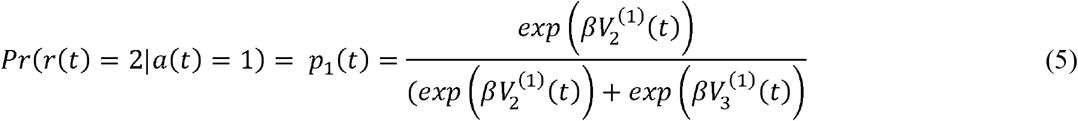

in which, *β* is the inverse temperature. The same deduction works for all players.

The Reciprocity model and the Alternating model each consider just one component when computing the values. Formally, for the Reciprocity model,

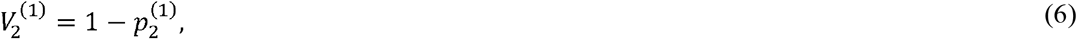

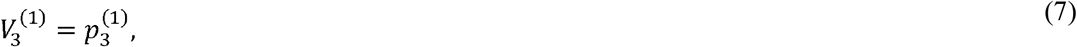

while for the Alternating model:

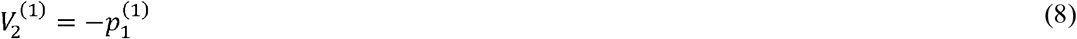

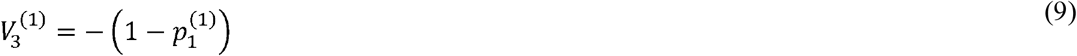

#### Guesses

As by the assumption of the naïve model, the player can only guess the choices of the Actors by their estimation of the Actors’ choosing tendency. Thus, in Experiment 3, the probability of the player *i(i ∈ N)*’s guessing about whether the player *j* would choose player *k* is 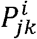

### The Mental Infer model

#### From the perspective of player 1

The naïve models only assume that the players track the actions of others and do not have a mental model for others. The Mental Infer model instead considers players’ mental model of others, which results in the fact that in *Player* 1’s mind, the action of player *i(i ∈ N)* would influence the sending probabilities of *Player j*(*j* ∈ *N* \ {1}). We assumed that in the Mental Infer model, the players model others behavior as follow the Alternating Reciprocity model.

To decipher this influence, we rewrote equation (5) for trial t + 1 as:

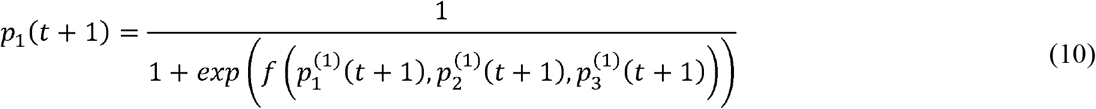

where *f* is the value difference, and we treated it as a function of the underlying probability estimates. We then insert equation (2) into equation (10), with the following substitution of variables:

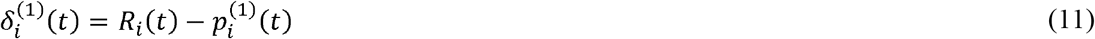

and rewrite *f* as a function of *η, f = f (η)*:

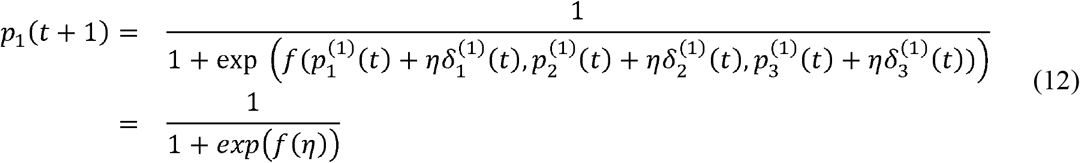

when *η* =0, equation (11) becomes *p*_1_*(t)*, so we can approximate Δ*p* =*p*_1_(*t* + 1) − *p*_1_*(t)* using Taylor Expansion near *η* = 0 as these two values are usually very small:

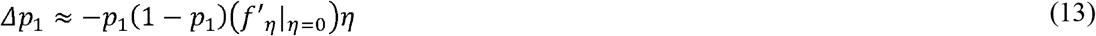

Note that we only take the first term of the Taylor expansion for this approximation. In 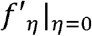 we take the partial derivative of *f* relative to *η* and take it’s value at *η* = 0. Then the only thing we need to do is to find 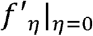. It’s easy to get these two values from equation (3) and (4).

For simplicity, we remove all the subscripts (1), and keep in mind that the equation is written in the view of player 1:

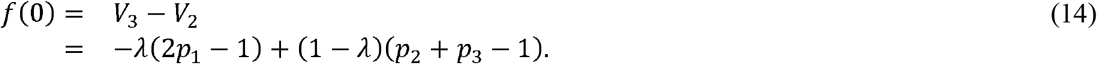

Then we have:

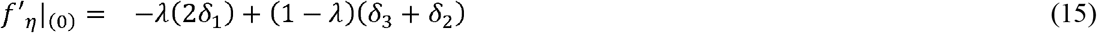

All *p, δ* are in the view of player 1 (i.e., we omitted the subscript (1) from the LHS).

#### Moving to the view of player 2

We now move to the viewpoint of player 2. Applying the same derivation as above we can obtain the update of player 2:

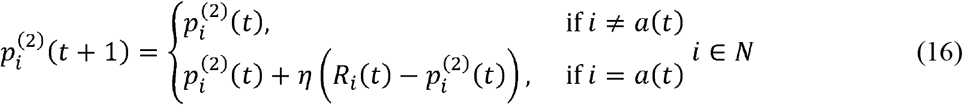

This equation says that when player *i* is acting, player 2 (hence all players) are updating their estimation of the player *i*’s probability of choosing player *ccw(i)*. The key to the Mental model is that the player thinks all other players act as an Alternating Reciprocity model and can use this mental representation to further adjust their estimation of *P* _*i, ccw(i)*_. The equation (15) provides such a possibility. Specifically, per the Mental Infer Model, player 2 knows:

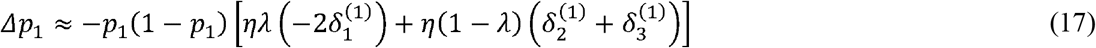

And s/he needs to substitute the estimation with his own estimation as:

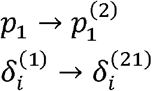

in which 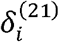 means in player 2’s mind, what is the player 1’s estimates of *δ*_*i*_, same explanation for other notations. We further assume in this stage, *player* 2 thinks layer 1 has the same estimation with him. This projection can be thought of as a higher level false belief documented in ToM task with high cognitive demand (Ereira et al., 2020) which makes the update possible:

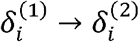

This leads to

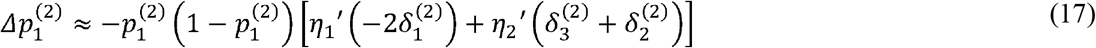

in which, *η*’ are learning rate. To further reduce the number of parameters, we let 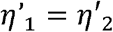, which guarantees the accuracy of the estimation of the parameters. This makes the equation (16) for *i* = 1 to:

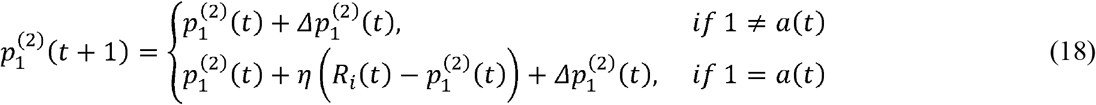

The above deduction applies to all 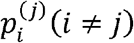 This completes our influence updating rules. One thing to note is that, for 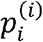, we do not assume Mental Model updating (the players do not use a Mental Model to predict their own decisions).

#### Guesses

As the Mental Infer model assumes the player represents other players as an Alternating Reciprocity model, the probability of the fact that player 2 guesses that player 1 would choose player 2 is given by:

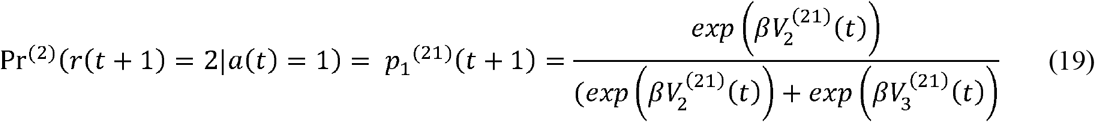

Note that, in this equation,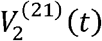 is a function of 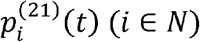 and player 1’s *λ*. By our assumption, 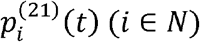 reduces to 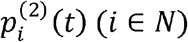 For player 1’s *λ*, we could treat it as a free parameter, but to simplify the model further, we assumed the players attributed their own weight to others. That is, in this equation, the *λ* is player 2’s own.

### Model Fitting And Simulation

All models were fit via Maximum Likelihood Estimation, employing a nonlinear optimization procedure using 100 random start points in the parameter space in order to find the best-fitting parameter values for each participant. We then computed the Akaike Information Criterion (AIC; (Akaike, 1974) to select the best-fitting models penalizing each model’s goodness-of-fit score by its complexity (i.e., number of free parameters).

To explore how the winning model (Mental Infer) beat other models, we simulated 100 artificial subjects for each participant with the best-fitted parameter value for each model, which resulted in one dataset for each model in each experiment. We then summarized the dataset of each model with the same analysis method as for the model-free analysis.

To verify if the free parameters of the winning model (Mental Inference) are recoverable, we simulated 100 artificial groups for each experiment with free parameters randomly chosen (uniformly) from their defined ranges. We then employed the same model-fitting procedure as described above to estimate these parameter values. We found that all parameters of this model could be recovered, while for the inverse temperature the correlation between simulated and recovered was relatively low but significant, a phenomenon found in many studies (Buergi et al., 2026; Zhang et al., 2025).

## Acknowledgements

This work was supported by the European Union (ERC Starting Grant, NEUROGROUP, 101041799; R.B.M.) and the Fundamental Research Funds for the Central Universities”(XJJSKYQD202501). Views and opinions expressed are however those of the authors only and do not necessarily reflect those of the European Union or the European Research Council Executive Agency. Neither the European Union nor the granting authority can be held responsible for them.

## Author Contributions

Conceptualization, S.Z. and R.B.M.; Methodology, S.Z. and R.B.M.; Formal Analysis, S.Z.; Investigation, S.Z. and H.W.; Writing, S.Z. and R.B.M.

